# Mitochondrial genomes and phylogenetic analysis of Central American weakly-electric fishes: *Apteronotus rostratus, Brachyhypopomus occidentalis* and *Sternopygus dariensis*

**DOI:** 10.1101/353565

**Authors:** Celestino Aguilar, Matthew J. Miller, Jose R. Loaiza, Rüdiger Krahe, Luis F. De León

## Abstract

Electric fishes are a diverse group of freshwater organisms with the ability to generate electric organ discharges (EODs) that are used for communication and electrolocation. Over 200 species have originated in South America, but of these, only a few have managed to colonize the Central American Isthmus. Here, we assembled two complete and one nearly complete mitochondrial genomes (mitogenomes) for three Central American Gymnotiformes: *Sternopygus dariensis, Brachyhypopomus occidentalis* and *Apteronotus rostratus*. We then explored the three species’ phylogenetic position in the context of South American electric fishes. Mitogenomes were organized in the standard fish mitogenome order, and presented sizes of 16,600, 16,540 and 15,940 base pairs (bp) (nearly complete) for *S. dariensis, B. occidentalis* and *A. rostratus*, respectively. We uncovered a novel 60 bp intergenic spacer (IGS) located between the *COII* and tRNA^Lys^ genes, which appears to be unique to the Apteronotidae. Furthermore, phylogenetic relationships supported the traditional monophyly of Gymnotiformes, with the three species positioned within their respective family. In addition, the genus *Apteronotus* was placed as the basal taxon of the order. Finally, we found high sequence divergence (13.3%) between our *B. occidentalis* specimen and a sequence previously reported in GenBank, suggesting that the prior mitogenome of *B. occidentalis* represents a different South American species that was misidentified. Indeed, phylogenetic analyses using *Cytochrome b* gene across the genus placed the previously reported individual within *B. bennetti*. Our study provides novel mitogenome resources that will advance our understanding of the diversity and phylogenetic history of Neotropical fishes.

## Introduction

Electric fishes (Teleostei, Gymnotiformes) are a highly diverse group of freshwater organisms that originated in South America (Albert, 2001). One of the defining features of these fishes is their ability to produce electric organ discharges (EODs) that are used for communication and electrolocation <u>(Moller 1995; Bullock et al. 2005)</u>. EODs are species-specific electric signals that can be divided into pulse-type and wave-type, depending on the shape and regularity of the discharge. In addition, there is evidence for reproductive character displacement in EOD waveform in this group (Crampton et al., 2011), which has over 200 currently described species.

Electric fishes are widely distributed in lowland freshwater habitats throughout South America (Albert and Crampton, 2005; Hulen et al., 2005). In Central America, however, only 6 species and five genera have been reported thus far, including *Apteronotus*, *Brachyhypopomus*, *Eigenmannia*, *Gymnotus* and *Sternopygus* (Alda et al., 2013; Reis et al., 2003). Despite the high diversity of Neotropical electric fishes, limited genomic resources are currently available for the group, particularly for Central American species. For instance, to date, only nine mitochondrial genomes of Gymnotiformes have been deposited in GenBank (Elbassiouny et al., 2016; Lavoué et al., 2012; Nakatani et al., 2011), but none of these mitogenomes belong to a Central American species. In the case of *B. occidentalis*, it is difficult to determine if the individual from South America previously reported by Lavoué et al. (2012) corresponds to the Central American species, particularly, because *B. occidentalis* presents a wide geographic distribution in South and Central America (Crampton et al., 2016b), and species-level divergence is likely to exist across the species’ range (Bermingham and Martin, 1998; Picq et al., 2014). This knowledge gap is important because the dynamic history of the Central American Isthmus has led to a complex evolutionary and phylogeographic history among electric and other primary freshwater fishes (Bermingham and Martin, 1998; Picq et al., 2014). Thus, generating molecular datasets – including complete mitogenomes – for Central American species of electric fishes is valuable to improve our understanding of diversification in Neotropical environments.

Here, we report for the first time full mitogenome sequences for three Central American weakly-electric fishes: two wave-type species – *Apteronotus rostratus* and *Sternopygus dariensis* – and the pulse-type *Brachyhypopomus occidentalis*. We also compile currently available mitogenomic data to assess the phylogenetic position of the three species within Gymnotiformes. In addition, we estimate genetic distances across complete mitochondrial genomes of three *Brachyhypopomus* individuals available in Genbank. Our study provides novel genomic resources that could facilitate further work on the conservation genetics, phylogenetics, and evolution of Central American Gymnotiformes as well as other freshwater fishes.

## 2. Materials and methods

### 2.1. Study site and sampling protocol

We collected three individuals from each of the following species: *A. rostratus, S. dariensis* and *B. occidentalis* in La Hoya stream, which flows into the Chucunaque River in the Darien Province, eastern Panama (N 8.2536, W -77.7189). Fish were detected using wire-electrodes connected to a mini-amplifier (Radio/Shack, Fort Worth, TX), and collected using a hand-net. Fish were euthanized with an overdose of eugenol (C_10_H_12_O_2_) derived from clove oil. Our collecting protocol was authorized by Ministerio de Ambiente (Mi Ambiente; permit number SE/A-100-14) and approved by the Institutional Animal Care and Use Committee (IACUC-16-001) at the Instituto de Investigaciones Científicas y Servicios de Alta Tecnología (INDICASAT AIP).

### 2.2. Sequencing

Mitochondrial genomes were obtained as the byproduct of Next Generation Sequencing (NGS) of Ultraconserved Elements (UCEs; Faircloth et al., 2012), as part of ongoing studies on the population genomics of the weakly-electric fish *B. occidentalis*. We prepared UCEs libraries following a standard protocol (available from http://ultraconserved.org) using the 500 loci Actinopterygii probe set (Actinopterygians 0.5Kv1; Faircloth et al., 2013). Libraries were sequenced using 300 bp paired-end Illumina MiSeq platform (Illumina, San Diego, CA) at the Smithsonian Tropical Research Institute (STRI) Naos Molecular Lab in Panama City, Panama.

### 2.3. Mitogenome assembly and annotation

We followed Aguilar et al. (2016) to generate mitogenomes from UCE sequencing reads. Briefly, we used Illumiprocessor (Faircloth, 2013), which employs Trimmomatic (Bolger et al., 2014) to clean and trim reads. We then assembled all reads using Trinity (Grabherr et al., 2011). Contigs larger than 15,000 bp were subjected to searches of sequence similarity using the BLAST algorithm to compare the query sequences with sequences from GenBank-NCBI (https://blast.ncbi.nlm.nih.gov/Blast.cgi) to confirm the presence of the mitochondrial genome. We annotated genes using the MitoFish and MitoAnnotator (Iwasaki et al., 2013) and also inspected alignments manually by comparing them with Genbank reference mitogenomes from *A. albifrons* (Accession no. AB054132), *S. arenatus* (Accession no. KX058571), *B. verdii* (*B. n.sp*. VERD – Accession no. KX058570) and *B. occidentalis* (Accession no. AP011570) in Geneious version 11.1.4 (http://www.geneious.com., Kearse et al., 2012).

Nucleotide base composition was also calculated in Geneious version 11.1.4 (http://www.geneious.com, Kearse et al., 2012). Strand asymmetry was calculated using the following formulas: AT-skew=(A−T)/(A+T) and GC-skew=(G−C)/(G+C), which allowed us to measure the nucleotide compositional difference between complete mitogenomes (Perna and Kocher, 1995).

### 2.4. Phylogenetic analysis

We used MAFFT (Katoh and Standley, 2013) to describe phylogenetic relationships among available mitogenomes of Gymnotiformes. For these analyses, we excluded the control region, and used the characiform *Astyanax paranae* as outgroup (Genbank accession no. KX609386). We tested for the best-fit model based on Akaike Information Criterion (AIC), corrected Akaike Information Criterion (AICc) and Bayesian Information Criterion (BIC), which identified the GTR+I+G model as best-fitting to our data. Bayesian inference (BI) and maximum-likelihood (ML) analysis were performed using the CIPRES Science Gateway v3.3 cluster (Miller et al., 2010). We performed two independent runs of MrBayes v.3.2.6 (Ronquist et al., 2012) using 8,000,000 Markov Chain Monte Carlo iterations (MCMC), with four simultaneous chains, and sampling every 1000 generations. Support for node and parameter estimates were derived from a majority rule consensus of the last 5,000 trees sampled after convergence. In addition, we generated a maximum likelihood phylogeny using RAxML (Stamatakis, 2014) with 1000 bootstrap replicates.

### 2.5. *Genetic divergence in* Brachyhypopomus *mitogenomes*

To assess genetic distances among the three *Brachyhypopomus* mitogenomes, we estimated the proportion of nucleotide differences (uncorrected p-distance; Nei and Kumar, 2000) for each protein coding gene separately. Standard errors were calculated using 500 bootstrap replicates in MEGA v7 (Kumar et al., 2016). We also estimated sequence divergence across species of *Brachyhypopomus* using *COI* (e.g., the barcoding gene; Hebert et al., 2003) data available in Genbank (Bermingham and Martin, 1998; Picq et al., 2014). Finally, to confirm the species identity of available *Brachyhypopomus* mitogenomes, we built a phylogenetic tree using a dataset of 83 *Cyt b* sequences representing 26 species reported in Crampton et al. (2016a).

## 3. Results and Discussion

### 3.1. Mitogenome structure

Minimum sequence cover for our three mitogenomes was 26X. The size of the complete mitogenomes was 16,600 bp for *S. dariensis* (Genbank accession no. MH399590) and 16,540 bp for *B. occidentalis* (Genbank accession no. MH399591), while the nearly complete mitochondrial genome of *A. rostratus* had 15,940 bp (Genbank accession no. MH399592). All three mitogenomes contained 2 ribosomal genes (12S and 16S), 22 tRNAs, 13 protein-coding genes (PCGs), as well as a control region (Figure 1). They also contained similar gene counts and organization as in other Gymnotiformes (Elbassiouny et al., 2016; Lavoué et al., 2012; Nakatani et al., 2011).

**Figure 1.**
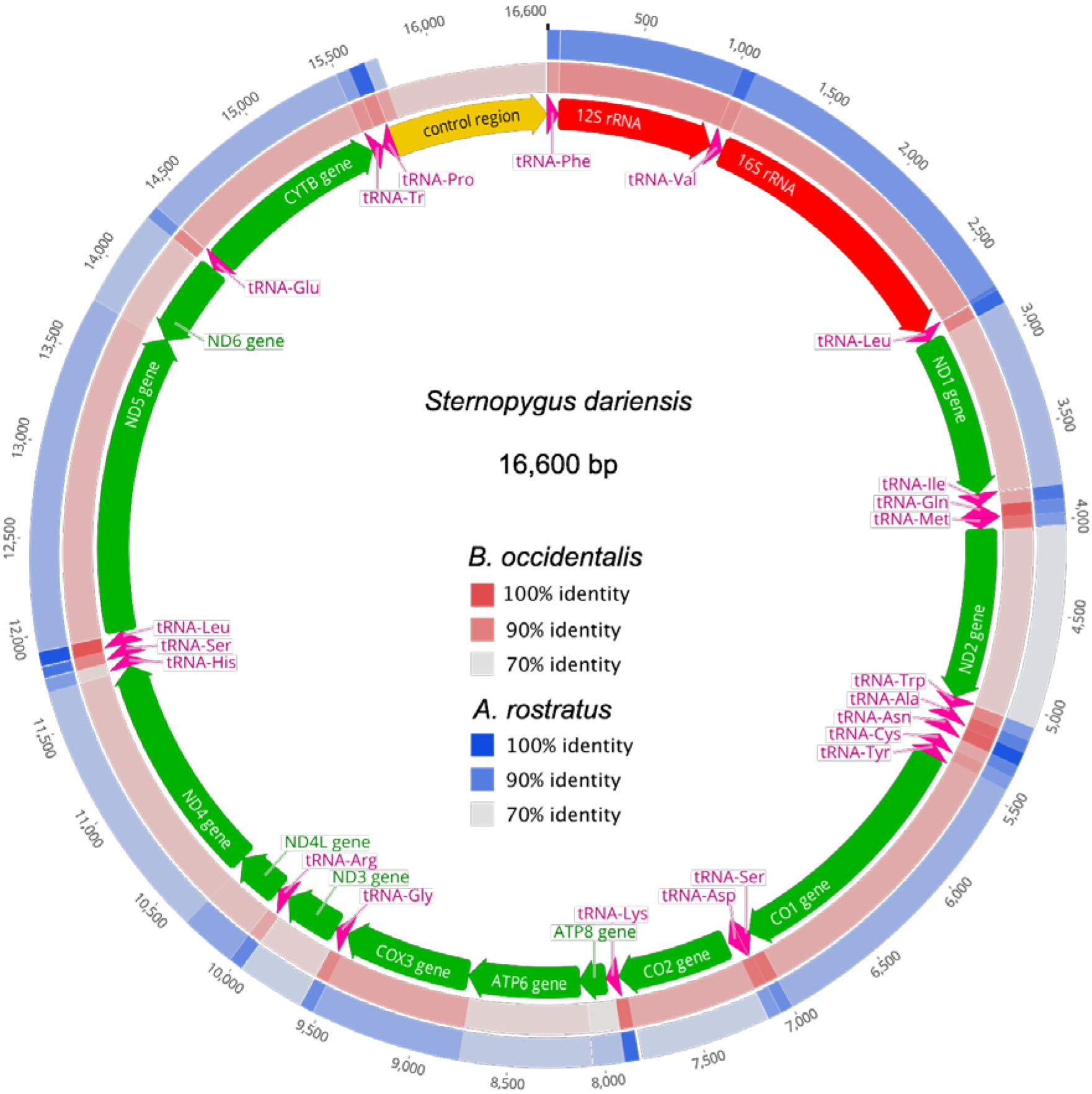
Comparison of the mitochondrial genome of *B. occidentalis* and *A. rostratus* against *S. dariensis*. Rings represent the mitogenome map of. *B. occidentalis* (outer ring), *A. rostratus* (middle), and *S. dariensis* (inner). The protein coding regions are labeled in green, tRNA genes are labeled in pink, rRNA genes are labeled in red and the putative control region is labeled in orange colors. The intensity of color of each of the two outer rings represents the proportion (70 to 100%) of conserved sequence at that region.

The nucleotide composition showed a strand bias consistent with the strand asymmetry observed in other fishes (Cheng et al. 2012; Hao et al. 2016). Specifically, the A + T content was 57.8%, 54.1% and 57.1% for *A. rostratus*, *B. occidentalis* and *S. dariensis*, respectively. The average AT-skew was 0.08, ranging from 0.05 in *B. occidentalis* to 0.09 in *S. dariensis*. The average GC-skew was −0.32, ranging from −0.34 in *S. dariensis* to −0.28 in *B. occidentalis*. The three mitogenomes also showed the typical structure of other Gymnotiformes (Elbassiouny et al., 2016). This included the 13 PCGs encompassing ~69% (11440 bp) of the total mitogenome; twelve of which were on the forward strand, while ND6 was on the reverse strand. Overall, we found ~3,791 codons, excluding stop codons, that were predicted for codon usage across the three mitogenomes. The start codon in *S. dariensis* and *B. occidentalis* was a typical ATN codon, but the start codon in the *COI* gene was GTG for all three species, which is consistent with other fish mitogenomes <u>(Satoh et al. 2016; Shi et al. 2016)</u>. However, *A. albifrons* mitogenome shows alternative start codons in three genes: GTG for *ATPase8* and *ND6*, and ACG for *ND4L*.

In fishes, the ACG start codon has been found particularly in the *A. albifrons ND4L* gene <u>(Satoh et al. 2016)</u>, but was also recently reported in the *ND1* gene of the perciform *Otolithes ruber* <u>(Guo et al. 2017)</u>. The three species shared the TAA stop codon in 3 PCGs *(ND1, ND4L and ND5)*, while AGA and AGG stop codons were present in *COI* (except in *B. occidentalis*, TAA) and *ND6* (except in *A. rostratus*, AGA), respectively. The remaining PCGs (*ND2, COII, ATPase6, COIII, ND3, ND4* and *Cyt B*) had TAG, TAA or the incomplete stop codons TA/T, which are presumably completed during post-transcriptional polyadenylation (Ojala et al., 1981). This pattern of stop codons is also common in other fishes (Kim et al., 2006; Nakatani et al., 2011). In addition, there were the typical 22 tRNAs predicted by Mitofish and tRNAscan, with a length ranging from 66 bp to 75 bp and including two *tRNA*^*Leu*^ and two *tRNA*^*Ser*^. The two rRNA genes were located between *tRNA*^*Phe*^ and *tRNA*^*Leu*^ and were separated by *tRNA*^*Val*^.

### 3.2. Non-coding regions, intergenic spacers and overlapping

We found small intergenic spacers (IGS) ranging in size from 1–60 bp, and totalling 58 bp in *S. dariensis*, 68 bp in *B. occidentalis* and 134 bp in *A. rostratus* (Table. 1). These IGS regions were mostly similar across species and represent a common feature of Gymnotiformes. One of these spacers, with a size of 29 to 31 bp, represents the origin of L-strand replication (OL), and is located between tRNA^*Asn*^ and tRNA^*Cys*^. We also found a large IGS of 955 bp that belongs to the Control Region (D-loop) in both *S. dariensis* and *B. occidentalis*. This spacer was only partially recovered in *A. rostratus*. We also observed 8 gene overlaps with a total of 31 bp; the two longest of which contained 10 bp (between *ATPase8* and *ATPase6*) and 7 bp (between *ND4L* and *ND4*) (Table 1).

**Table 1a.**
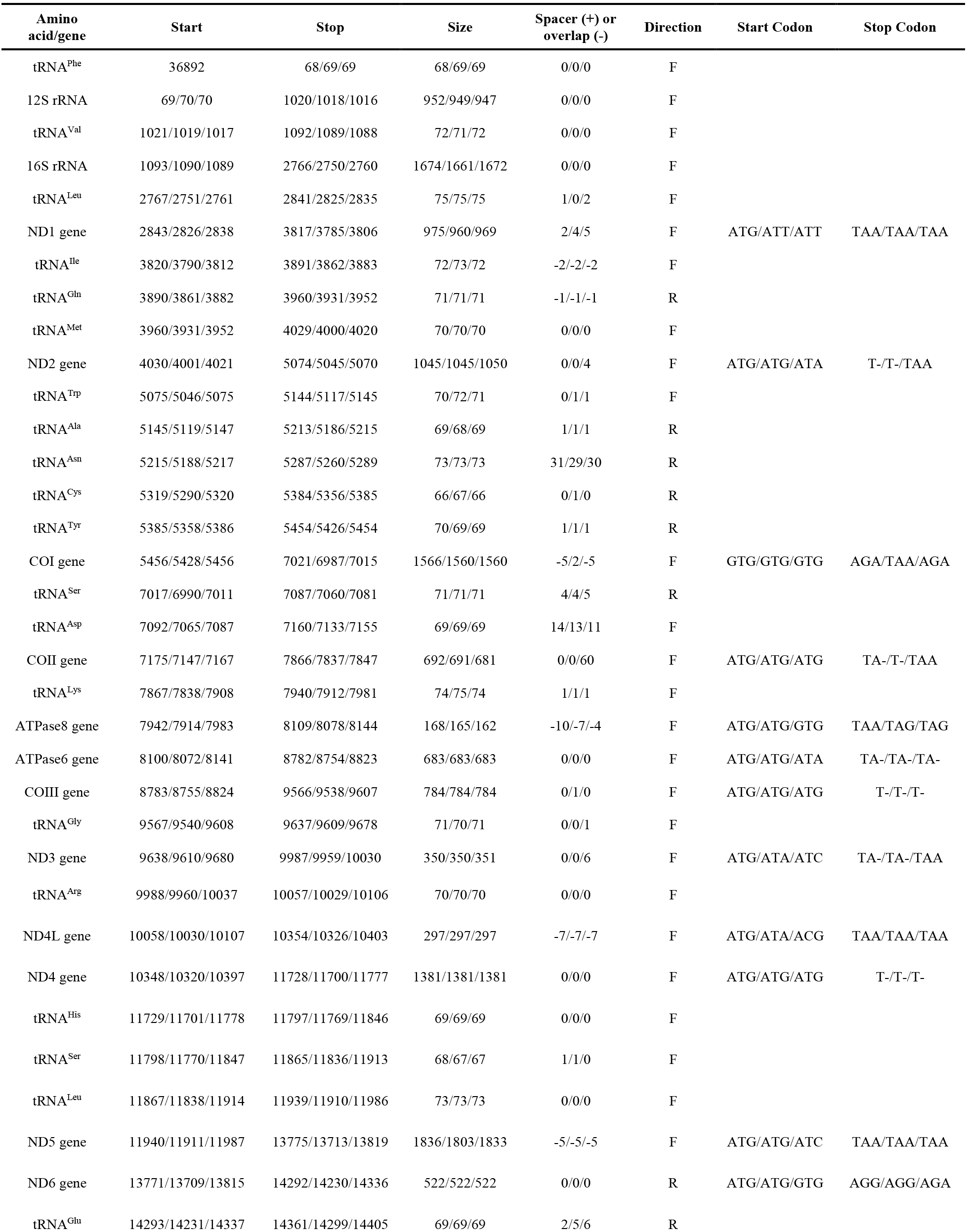
Characteristics of the mitochondrial genomes of Central American electric fishes. The table shows the mitogenome structure of three species in the following order: *S. dariensis, B. occidentalis* and *A. rostratus*.

**Table 1b.**
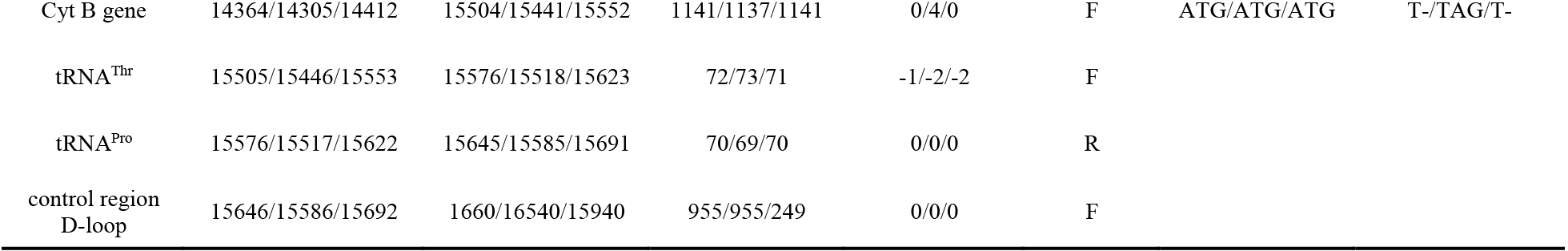

### 3.3. Novel COII/tRNA-Lys intergenic spacer in Apteronotus

We uncovered a 60 bp IGS between *COII* and tRNA^Lys^ in both *A. rostratus* (from present study) and *A. albifrons* (from GenBank; AB054132). However, this IGS was not found in any of the other available gymnotiform mitogenomes (Figure 2). This spacer showed clear similarity between the two *Apteronotus* mitogenomes that were sequenced independently, which suggests that this IGS represents a unique feature of the genus *Apteronotus*. To our knowledge, this is the first report of a long IGS occurring between the genes *COII* and tRNA^Lys^ in the order Gymnotiformes. In other fishes, the presence of unique IGS has been reported between the genes tRNA^*Thr*^ and tRNA^*Pro*^ in Gadiformes (Bakke et al., 1999; Jørgensen et al., 2014), including walleye pollock, *Theragra chalcogramma* (Poulsen et al., 2013), whiting *Merlangius merlangus* and haddock *Melanogrammus aeglefinushiting* (Roques et al., 2006).

**Figure 2.**
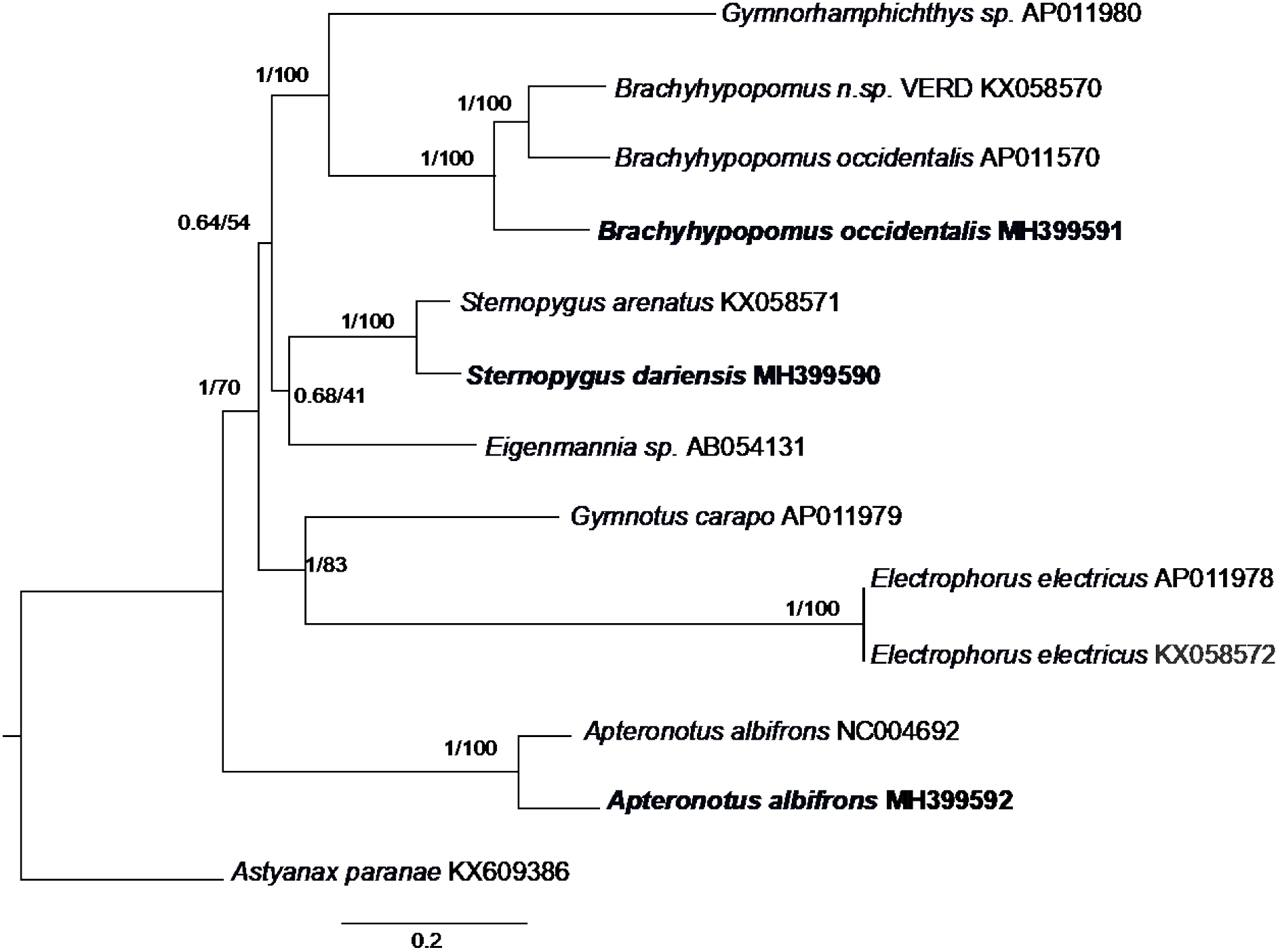
Phylogenetic relationships among Gymnotiformes based on MrBayes and RAxML. The phylogeny represents the best-scoring maximum likelihood tree based on complete mitogenomes (excluding the D-Loop). The first number at each node is Bayesian posterior probability and the second number is bootstrap probability of ML analyses. The scale bar indicates relative branch lengths.

Currently, the origin of these IGS and their apparent absence in other Gymnotiformes is not well understood. If we accept a basal phylogenetic position of *Apteronotus* (Figure 3), one possibility is that purifying selection on non-coding regions (Rand, 1993) led to reduction in mitogenome size during the evolutionary history of Gymnotiformes. However, further work is necessary to determine the biological implications of these IGS in Gymnotiformes. Overall, we suggest that comparative studies of this unique mitogenomic feature across species could help elucidate the phylogenetic history of the group. In addition, further work should explore the use of these IGS as genetic markers for the genus Apteronotus or the entire Apteronotidae.

**Figure 3.**
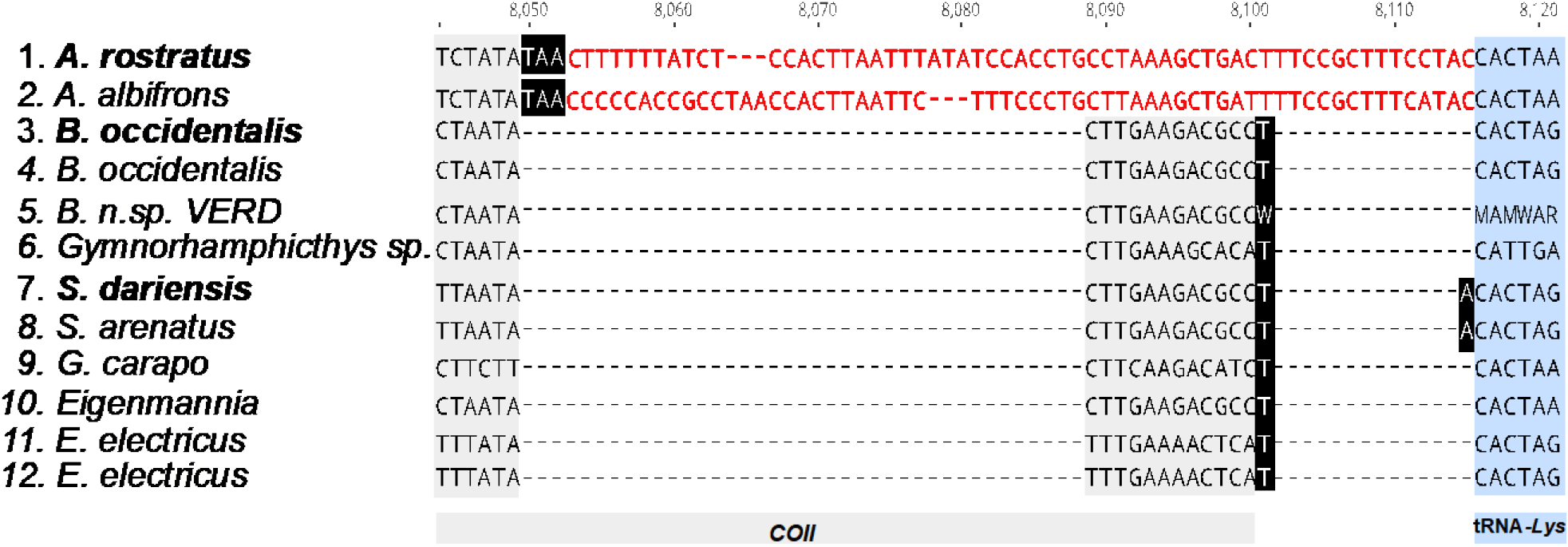
Alignment of partial mitogenome sequences of Gymnotiformes species. The 60 bp intergenic spacer, colored in red, is observed in the mitogenome of *A. rostratus* and *A. albifrons*. Individuals in bold represent sequences from the present study. Genes in color are encoded on the light strand, *COII* (grey) and *tRNA*^*Lys*^ (blue). Complete and partial stop codons are shown in black. Gaps are represented by dashes.

### 3.4. Phylogenetic and taxonomic implications

*Sternopygus dariensis* and *S. arenatus*, *A. rostratus* and *A. albifrons* as well as *B. occidentalis*, *B. verdii* (*B*. n.sp. VERD - Genbank KX058570) and *B. occidentalis* (AP011570) showed a monophyletic relationship with respect to each genus (Figure 2), confirming the phylogenetic position of Central American electric fishes within Gymnotiformes as a whole, consistent with Elbassiouny et al. (2016) and Tagliacollo et al. (2016). Our phylogenetic analysis recovered the monophyly of the order Gymnotiformes, and placed *Apteronotus* at the base of the order, in agreement with a recent mitogenomic study (Elbassiouny et al., 2016), but in contrast to the conclusions of Tagliacollo et al. (2016). Whereas the genetic results of the latter study were inconclusive with respect to the position of the apteronotids, their morphology-based tree identified Apteronotidae as a derived group within the Sinusoidea (Sternopygidae and Apteronotidae).

We found over 13% sequence divergence between our complete *B. occidentalis* mitogenome, and the one conspecific mitogenome available in Genbank. At individual genes, our analysis of p-distances revealed values ranging from 12.2% in *COII* to 19.9% in *ND4L*. For protein coding genes, average divergence was 15.6% (Table 2). We believe that our mitogenome represents the correct sequence for *B. occidentalis* given the 99.8% similarity between our sequence and previously sequenced individuals of *B. occidentalis* from Panama (Picq et al., 2014), but only 90.6% similarity with the *B. occidentalis* mitogenome reported by Lavoué et al. (2012; Accession no. AP011570). Indeed, our subsequent phylogenetic analysis of the genus *Brachyhypopomus* using *Cyt b* data from Crampton et al. (2016a) placed that individual within the *B. bennetti* clade from South America rather than with *B. occidentalis* of Central America (Figure S1).

**Table 2.**
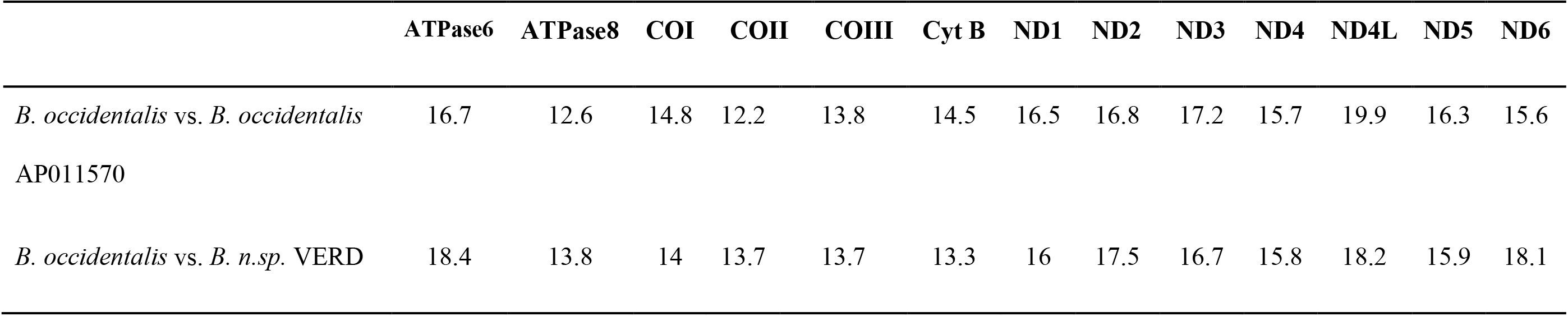
Pairwise genetic distance between specimens of *Brachyhypopomus*. The data represent percentage of uncorrected ‘p’ genetic distance between *B. occidentalis* vs. *B. occidentalis* (AP011570) and *B. occidentalis* vs. *B. n.sp*. VERD, based on 13 protein-coding genes (PCGs).

|  | ATPase6 | ATPase8 | COI | COII | COIII | Cyt B | ND1 | ND2 | ND3 | ND4 | ND4L | ND5 | ND6 |
| --- | --- | --- | --- | --- | --- | --- | --- | --- | --- | --- | --- | --- | --- |
| *B. occidentalis* vs. *B. occidentalis*<br>AP011570 | 16.7 | 12.6 | 14.8 | 12.2 | 13.8 | 14.5 | 16.5 | 16.8 | 17.2 | 15.7 | 19.9 | 16.3 | 15.6 |
| *B. occidentalis* vs. *B. n.sp.* VERD | 18.4 | 13.8 | 14 | 13.7 | 13.7 | 13.3 | 16 | 17.5 | 16.7 | 15.8 | 18.2 | 15.9 | 18.1 |

Overall, our study expands our understanding of the evolution and structure of mitochondrial genomes in Central American freshwater fishes. In addition, it generates novel molecular data that can be used to solve the taxonomic status as well as the phylogenetic history of Neotropical electric fishes.

*ATPase6* and *ATPase8*, ATPase subunit 6 and 8 genes; *Cyt B*, cytochrome b gene; *COI-III*, cytochrome oxidase subunits I-III genes; NCR, non-coding region; *ND1-6* and *ND4L*, NADH dehydrogenase subunits 1-6 and 4L genes; ML, maximum likelihood; BI, Bayesian inference; *rRNA*, ribosomal RNA; *16S* and *12S*, large and small subunits of ribosomal RNA genes; *tRNA*, transfer RNA; PCG, protein coding gene; mitogenome, mitochondrial genome; *Ala*, alanine; *Arg*, arginine; *Asn*, asparagine; *Aps*, aspartic acid; Cys, cysteine; *Gln*, glutamine; *Glu*, glutamic acid; *Gly*, glycine; *His*, histidine; *Ile*, isoleucine; *Leu*, leucine; *Lys*, lysine; *Met*, methionine; *Phe*, phenylalanine; *Pro*, proline; *Ser*, serine; *Thr*, threonine; *Trp*, tryptophan; *Tyr*, tyrosine; *Val*, valine.

## Highlights

1. The mitochondrial genomes of three Central American electric fishes, *Apteronotus rostratus, Brachyhypopomus occidentalis* and *Sternopygus dariensis*, are sequenced and characterized.
2. The presence of a novel 60 bp intergenic spacer located between *COII* and tRNA^Lys^ is reported for the first time in Gymnotiformes and may represent a unique feature of the *Apteronotus* mitogenome.
3. Phylogenetic analyses support the position of Central American *A. rostratus, B. occidentalis* and *S. dariensis* within monophyletic Gymnotiformes.
4. Genetic divergence and phylogenetic analyses indicate that the mitogenome of *B. occidentalis* (Genbank AP011570) reported previously belongs to *B. bennetti* from South America.

## Supplemental material

**Figure S1.**
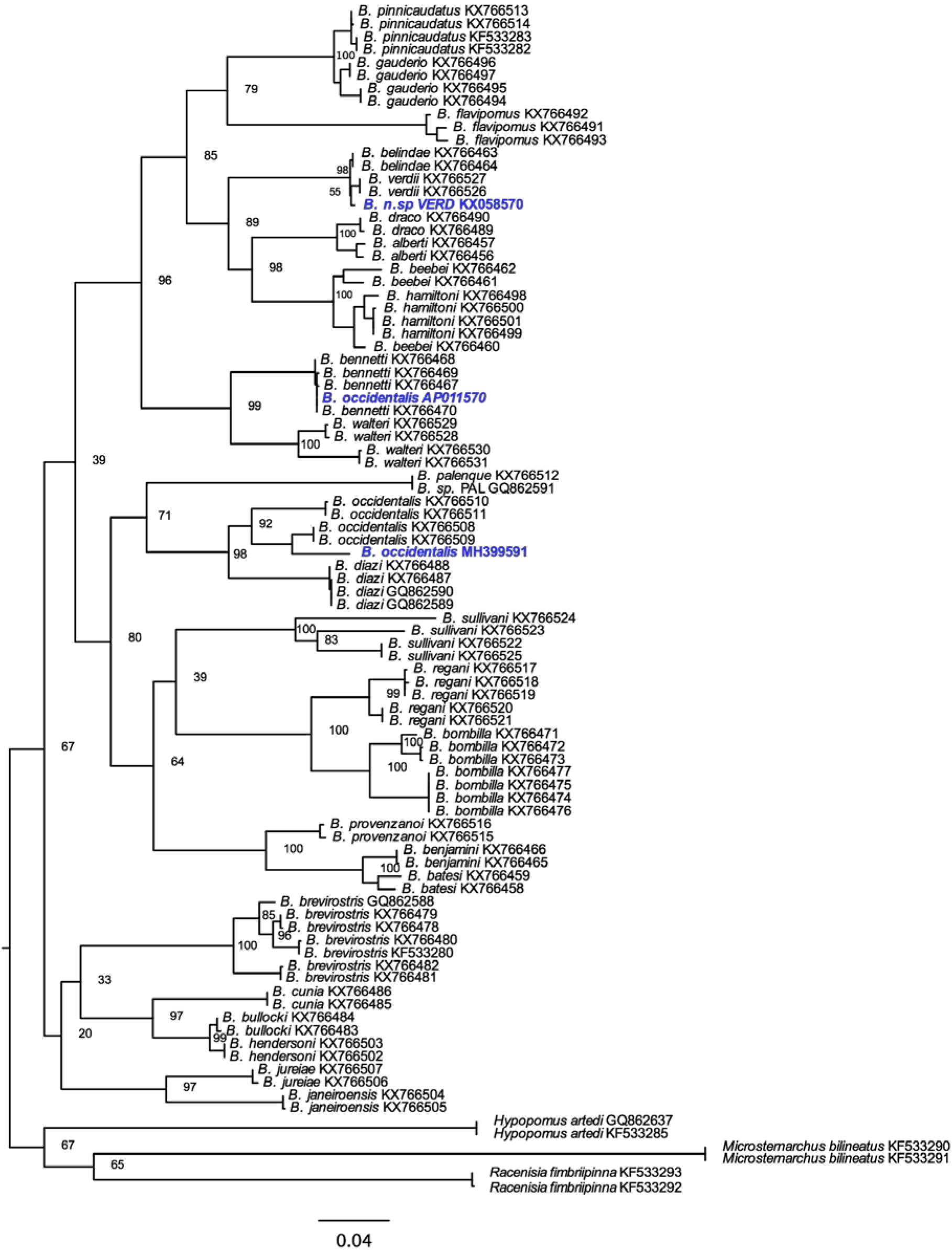
RAxML phylogenetic tree of 26 *Brachyhypopomus* species based on the *Cyt b* gene. Individuals in blue represent mitogenomes reported here (Genbank MH399591) and two additional sequences obtained from GenBank (AP011570, KX058570).

